# Prior epigenetic status predicts future susceptibility to seizures in mice

**DOI:** 10.1101/2025.03.20.644199

**Authors:** Benjamin D. Boros, Mariam A. Gachechiladze, Juanru Guo, Dylan A. Galloway, Shayna M. Mueller, Mark Shabsovich, Allen Yen, Alexander J. Cammack, Tao Shen, Robi D. Mitra, Joseph D. Dougherty, Timothy M. Miller

## Abstract

Wide variation of responses to identical stimuli presented to genetically inbred mice suggests the hypothesis that stochastic epigenetic variation during neurodevelopment can mediate such phenotypic differences. However , this hypothesis is largely untested since capturing pre-existing molecular states requires non-destructive, longitudinal recording. Therefore, we tested the potential of Calling Cards (CC) to record transient neuronal enhancer activity during postnatal development, and thereby associate epigenetic variation with a subsequent phenotypic presentation – degree of seizure response to the pro-convulsant pentylenetetrazol. We show that recorded differences in epigenetics at 243 loci predict a severe vs. mild response, and that these are enriched near genes associated with human epilepsy. We also validated pharmacologically a seizure -modifying role for two novel genes, *Htr1f* and *Let7c*. This proof-of-principle supports using CC broadly to discover predisposition loci for other neuropsychiatric traits and behaviors. Finally, as, human disease is also influenced by non-inherited factors, similar epigenetic predispositions are possible in humans.

## INTRODUCTION

Neuropsychiatric disease risk, while heavily impacted by genetic makeup, is also influenced by non-genetic factors during development. For example, epigenetic states early in life may be established in response to factors such as environmental exposures (e.g., head traumas) or psychosocial background. Thus, inter-individual differences could be attributed partially to naturally-occurring variation in epigenetic states across the population. This epigenetic variation could contribute to brain disorders like epilepsy, which can have variable presentation even between identical twins (Berkovic et al., 1998; Kjeldsen et al., 2005; Krenn et al., 2019). However, a substantial challenge in studying epigenetic risk factors is disentangling pre-existing molecular states from those occurring as a consequence of an experimental manipulation or the disease itself. Overcoming this challenge would represent an important milestone, as connecting non-inherited variation to stochastic differences in pre-existing epigenetic states could help identify novel mediators of seizure propensity or other naturally variable neuropsychiatric disease traits.

Epilepsy affects 46 million individuals worldwide and is characterized by unprovoked, recurrent seizures that can be highly variable in onset, frequency, duration, or intensity (Beghi et al., 2019; Devinsky et al., 2018). Human genome-wide association studies (GWAS) have identified many inherited risk factors for epilepsy. Many monogenic forms are causally linked to loss- or gain-of-function mutations in proteins influencing membrane excitability, such as *SCN1A* or *GABRA1* (M.-W. Zhang et al., 2024). Furthermore, other GWAS implicated genes like *BCL11A* or *RORB* are transcription factors that influence neuron development and identity, rather than membrane excitability (Abou-Khalil et al., 2018; Stevelink et al., 2023), and many gene variants that cause neurodevelopmental disorders or intellectual disability (ID) broadly also exhibit comorbid epilepsy (Zacher et al., 2021). Nevertheless, even among individuals with the same mutation, seizure frequency and intensity can vary substantially, suggesting other processes influence risk. Indeed, it is well-established that non-genetic factors such as traumatic brain injury or stroke also increase risk for developing seizures, and in some at-risk individuals, stress, alcohol consumption, or sleep deprivation can trigger an episode (Wassenaar et al., 2014). These lines of evidence argue that non-genetic risk factors arising stochastically during typical neurodevelopment, or that are uniquely acquired, may also influence seizure risk.

Variability in neuropsychiatric phenotypes influenced by non-genetic factors also exists in animals. For example, mouse models of acute seizure (e.g., induction with GABA-A receptor antagonist pentylenetetrazol; PTZ), result in a range of seizure responses even in effectively isogenic mouse strains (Van Erum et al., 2019). Specifically, while most mice experience generalized tonic-clonic activity, some only exhibit mild freezing while others proceed to death. We hypothesize that this natural variability in presentation might be due to stochastic variation in pre-existing epigenetic states established during brain development. We therefore posited that PTZ induction in naive mice might represent a model system to examine 1) whether such pre-existing molecular states can influence subsequent phenotype or presentation, and 2) if so, to identify naturally-occuring susceptibility factors from such molecular states.

However, most methods of assessing molecular state, such as RNA-seq or ChIP-seq, are destructive, and can therefore only provide a snapshot of the molecular state at the time of sacrifice. They cannot, therefore, be used to connect pre-existing epigenetic or molecular differences to behavioral outcomes (Johnson et al., 2007).Calling Cards (CC) is a transposon-transposase based technology that can create a permanent record of one such molecular state: transient transcription factor-DNA interactions (Cammack et al., 2020; Wang et al., 2012; Yen et al., 2023). The crux of CC is a self-reporting transposon that is directed to a genomic locus by the transposase *hyperPiggyBac*, which has a natural affinity for BRD4 binding sites in the genome. BRD4 is a histone acetylation reader that preferentially binds at active enhancers and promoters; increased levels of BRD4 binding often predict increased gene expression (Devaiah et al., 2016; Kanno et al., 2014). The transposase-DNA interaction results in a transposon insertion at that locus, serving as a permanent record of binding that can be read out through targeted sequencing after sacrifice. Unlike epigenetic profiling tools like ChIP-seq or ATAC-seq, CC can record enhancer activity across extended time windows (i.e., across development prior to PTZ injection). Because CC insertion and subsequent transposon reporter expression requires relatively long time windows (24-48 hrs), minimal recording of molecular state occurs in the few minutes between PTZ injection and seizure (in contrast to rapid RNA induction of immediate early genes like c-fos). Therefore, CC could circumvent the chief challenge of disentangling pre-existing molecular states from those occurring as a consequence of seizure induction. If so, then CC could be applied broadly to understanding variability in many other neurological diseases or behaviors beyond acute seizure induction (**Fig 1**).

**Figure 1.**
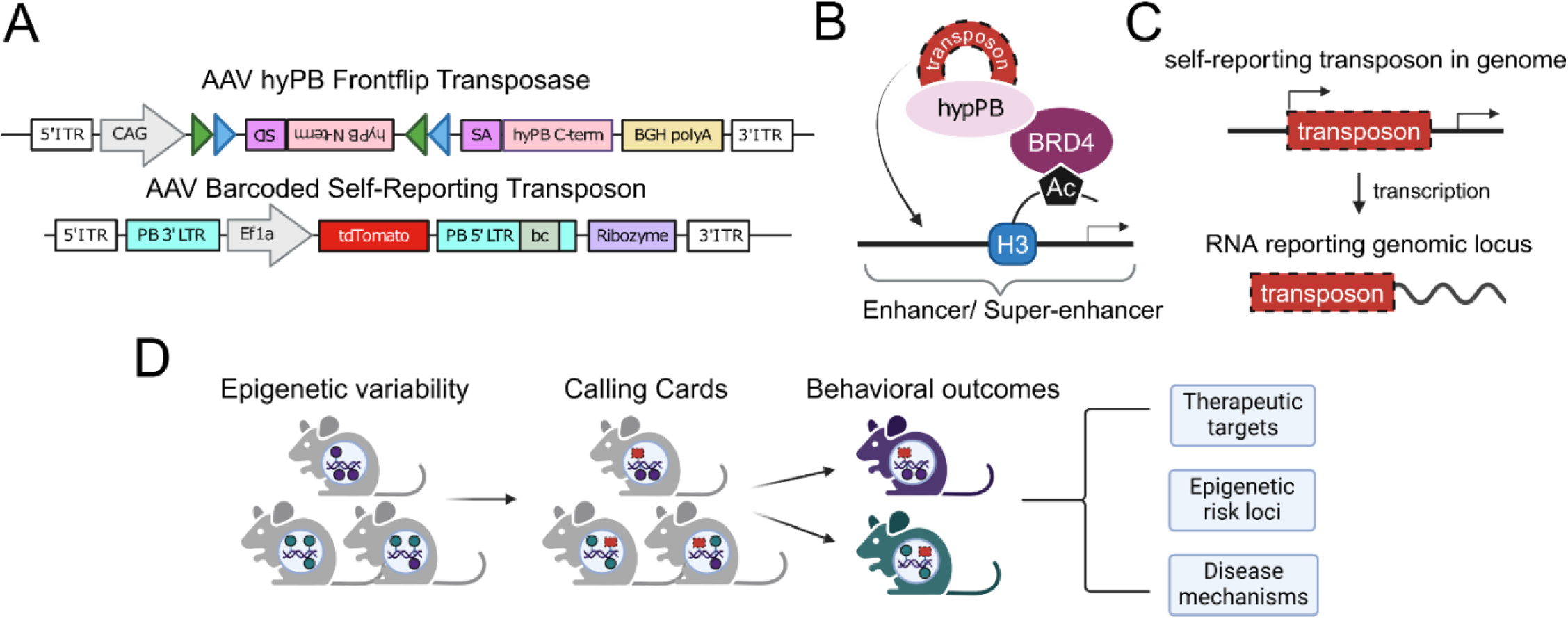
Paradigm for using Calling Cards to identify genomic loci associated with behavioral outcomes. (A) Frontflip hyPB transposase (top) and barcoded self-reporting transposon (bottom) constructs. (B) Unfused hyPB (hyper Piggyback) transposase naturally interacts with BRD4 and directs donor transposons to enhancer sites, as marked by histone 3 (H3) acetylation (Ac). (C) A self-reporting transposon (red box) is inserted into the genome that reports its genomic locus via transcription. (D) Approach to use Calling Cards to connect pre-existing epigenetic states with behavioral variability in order to identify novel disease mechanisms, therapeutic targets, and epigenetic risk loci. Insertion of CC transposons are denoted by the presence of red transposon boxes into the genome.

Here, we examined whether such a pre-recorded epigenetic status can predict future seizure susceptibility. We implemented neuron-specific CCs immediately after birth to record epigenetic activity through one month of age. We then induced acute seizures and sacrificed animals shortly thereafter. Next, we stratified mice based on seizure responses and tested for differential CC transposon insertions between severe vs. mild seizure responders. We identified numerous loci with pre-existing epigenetic differences between these groups. Further, putative enhancer sites mapped near known epilepsy risk genes, and susceptibility loci were enriched for genes influencing neurodevelopmental maturity. Finally, we implemented pharmacological methods to support the identification of two novel seizure-modifying genes from our dataset. This work supports the hypothesis that pre-existing epigenetic states that affect novel or established epilepsy pathways can influence seizure susceptibility. Our study thus provides a template for examining antecedent epigenetic states for various other neuropsychiatric diseases and traits.

## MATERIALS AND METHODS

### Animals

All practices and procedures using mice were approved by the Washington University in St. Louis Institutional Animal Care and Use Committee in accordance with National Institutes of Health (NIH) guidelines. All mice used in this study were bred and maintained in the Washington University in St. Louis vivarium, where they were kept on a 12-hour light/dark cycle and had unlimited access to food and water. All mice were on a C57BL/6J background from Jackson Laboratory (catalog: 000664), and experiments were balanced for sex. “Syn1-Cre” mice used for implementation of Calling Cards were ordered from Jackson Laboratory: B6.Cg-Tg(Syn1-cre)671Jxm/J (catalog: 003966) and were inbred with C57BL/6J. Non-transgenic mice were used in drug studies and were euthanized in their home cage using carbon dioxide.

### Calling Cards Constructs, Virus Production, and Injections

Transposase and donor transposon plasmids were packaged into AAV9 capsids by the Hope Center Viral Vectors Core at Washington University School of Medicine, as described (Cammack et al., 2020). For the donor transposons, we used a barcoded pool of Self-Reporting Transposons to increase our ability to detect unique insertions, as described (Lalli et al., 2022). SRTs report their location via RNA transcription and thus enhance sensitivity and recovery (Moudgil et al., 2020) (Addgene: 1000000213), and were co-injected with the ‘Front Flip’ Cre-dependent Transposase, which cleanly drives activity only in Cre-expressing cells (ref Cammack). Titers for each virus ranged between 6.8 x 10^13 viral genomes/ml. A 1:1 mixture by volume of transposase and donor transposon viruses was intracranially injected into the cortex of P0-1 Syn1-Cre male and female mice (two sites per hemisphere; 1 μL of viral mix per site). Prior to injections, pregnant females were monitored daily, and newborn pups were injected within 24 hours. CC injected pups were not weaned prior to sacrifice at P28.

### Seizure Induction for Calling Cards and Evaluation

Mice were injected intraperitoneally with 65 mg/kg pentylenetetrazol (PTZ) (Sigma catalog: P6500-25G) and placed in a standard home cage, where they were evaluated for seizure activity for 15 minutes while also being recorded with a video camera. Immediately afterwards, mice were euthanized and decapitated. Cortices were extracted, and each hemi-cortex was dissected into three parts and flash frozen in liquid nitrogen. For CC libraries, Syn1-Cre animals were used at P28; for validation drug studies, non-transgenic mice were used at either P28 or at 3 months of age.

A modified Racine scale was used to score seizure responses during the 15 minutes period of observation (Van Erum et al., 2019) (**Table 1**.) For the purposes of this study, we assigned scores of 1-4 as “mild” responses; score 5 as “moderate”; and scores 6-8 as “severe.” The time at which each mouse achieved a score threshold was marked, and the latency from PTZ injection was calculated for time to “first response”, “score 3” (tail pop), or “score 5” (generalized tonic clonic activity). Comparisons of seizure latencies between groups were performed using only mice that achieved at least that score.

**Table 1.**
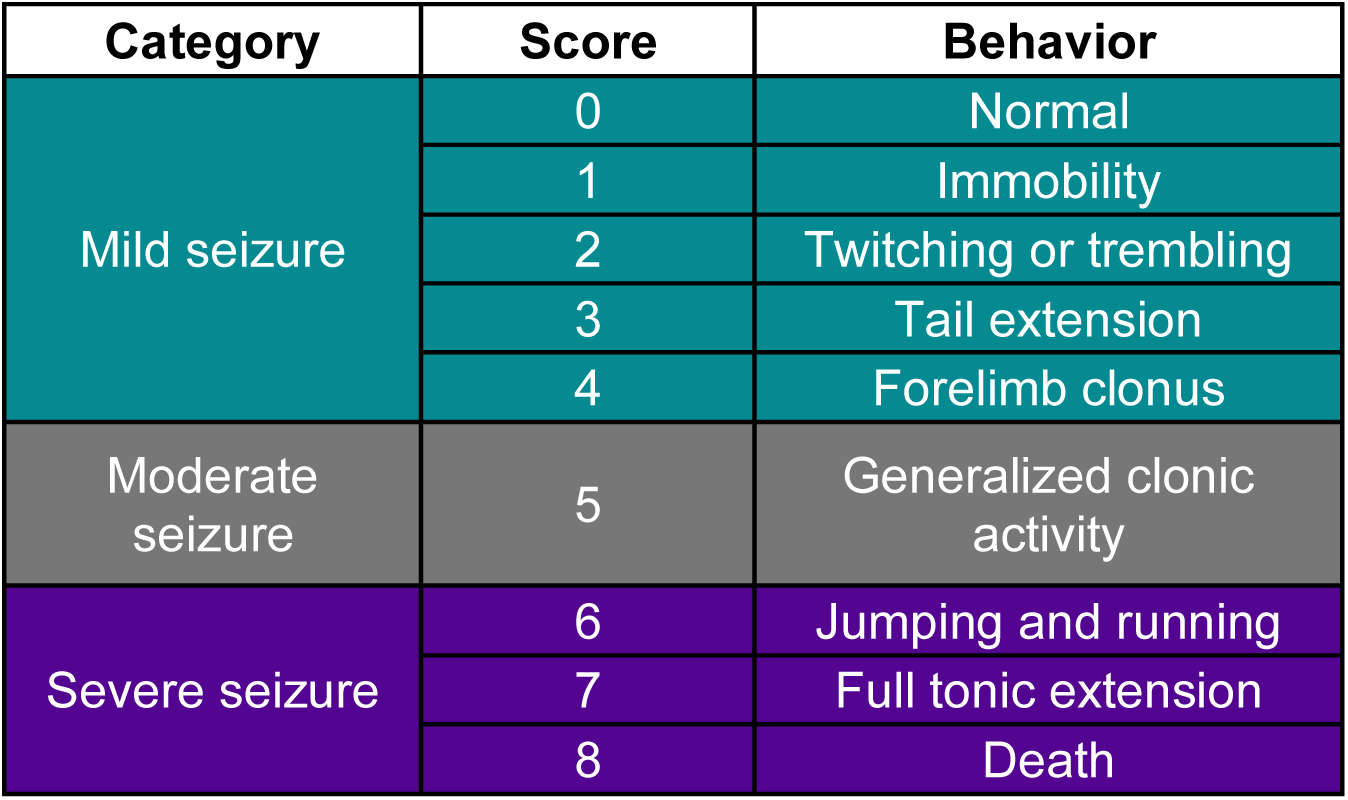
Scoring of seizure responses and division into mild, moderate, and severe seizures. Using a modified Racine scale, responses to PTZ are scored from 0 to 8, spanning “normal” behavior to “generalized clonic activity” to “death.”

| Category | Score | Behavior |
| --- | --- | --- |
| Mild seizure | 0 | Normal |
|  | 1 | Immobility |
|  | 2 | Twitching or trembling |
|  | 3 | Tail extension |
|  | 4 | Forelimb clonus |
| Moderate seizure | 5 | Generalized clonic activity |
| Severe seizure | 6 | Jumping and running |
|  | 7 | Full tonic extension |
|  | 8 | Death |

### Calling Cards Library Preparation and Sequencing

Calling Card protocol proceeded largely as described (Yen et al., 2023), with multiple cortical pieces (up to six per animal) processed separately to further increase our ability to detect unique insertions even beyond the number of available barcodes. Data were then pooled by animals for subsequent analysis. Briefly, RNA was extracted from cortical tissues according to the manufacturer’s instructions (Zymo RNA Clean & Concentrator-5 Kit and QIAGEN RNEasy Plus Mini Kit). SRT sequencing libraries were prepared from these RNA samples as previously described (Yen et al., 2023)and sequenced on the NovaSeq X Plus platform (Illumina).

Raw reads were processed into QBED files containing insertions at TTAA sites using the nf-core/callingcards nextflow pipeline (Yen et al., 2023) set with the default mammalian parameters for mouse. Reads aligning to the same TTAA site with separate barcodes, or from spatially separate cortical pieces, were considered unique insertions. After filtering for insertions with read depth >1, all unique insertions were considered equally, regardless of read depth.

### Calling Cards Peak Calling and Annotation

Significantly enriched insertion peak calling was performed with the pycallingcards module (Guo et al., 2024). Insertion profiles for all animals within each severity category were merged to create mild and severe profiles. For quality control, correlation coefficients were calculated on insertion profiles by animal within each category using the Pearson test. Animals with average correlation coefficients above > 0.91 were used for downstream analysis, resulting in 10 animals per group. Peaks were called on the mild and severe profiles separately using the MACCs method, then merged using pybedtools for a final set of peaks. The following parameters were used for the peak calling: reference = "mm10", pvalue_adj_cutoff = 0, window_size=2000, step_size=800, lam_win_size = 1000000, pseudocounts = 1.

Peak annotation was performed using bedtools(Quinlan & Hall, 2010) and HOMER(Heinz et al., 2010). Because enhancers usually exert their effects on nearby genes (Sun et al., 2019; Symmons et al., 2014), genes were assigned to peaks based on proximity. We used bedtools to find up to five of the closest genes within one megabase of the peak, filtering out “Gm” and “Rik” genes that have not been assigned names. Genomic features of the peaks were identified by HOMER.

For the label shuffling analysis, animal profiles were shuffled into two groups in 20 random arrangements such that each group had equal numbers of mild and severe animals and roughly equal numbers of insertions. Downstream peak calling and differential analysis proceeded as described here.

### Histone Modification Enrichment Analysis

Qualitative enrichment analysis of histone modifications at CC peaks was performed using deepTools(Ramírez et al., 2016). The following P0 mouse forebrain ChIP-Seq(He et al., 2020) datasets from ENCODE(Dunham et al., 2012) were used: H3K27ac (ENCSR094TTT), H3K4me1 (ENCSR465PLB), and H3K27me3 (ENCSR070MOK).

### Differential Peak Analysis

For the main differential peak analysis, we compared the number of insertions falling within each peak between mild and severe groups (or between shuffled groups) using Fisher’s exact test implemented in pycallingcards (Guo et al., 2024). Peaks with adjusted p-values < 0.05 after Benjamini-Hochberg correction were considered differential peaks. In addition, as a post-hoc we confirmed the correlation between insertion counts per million at each of the significant peaks and seizure score using the Spearman test, with Benjamini-Hochberg correction.

### Gene Ontology Enrichment Analysis

Gene ontology was performed using Panther classification system to identify enrichment in “biological” processes. Genes associated with mild responders or with severe responders were evaluated against a background of all genes annotated with CC peaks (16,240 genes). Processes achieving p<0.05 after Benjamini-Hochberg FDR correction are displayed.

### Gene Overlap Significance Testing

The significance of overlap between gene sets was statistically evaluated. The observed overlap between significant gene sets (mild, severe, both) and disease-related genes was compared to an expected distribution. This distribution was generated through 10,000 random permutations. In each permutation, genes were randomly sampled without replacement from the list of all genes annotated to CC peaks, with the sample size matched to each significant gene set being tested. The overlap between these random gene sets and the disease gene list was calculated. An empirical p-value was determined by dividing the number of permutations with equal or greater overlap than the observed value by the total number of permutations. Note the random gene lists and the significant sets are both drawn from peaks found in neurons, and thus this controls for any general neuron-specific bias to any disease.

The following disease gene datasets were used for the overlap analysis: monogenic epilepsy (Prevention Genetics Epilepsy and Seizure Panel; https://www.preventiongenetics.com/testInfo?val=Epilepsy-and-Seizure-Panel), epilepsy-related (Prevention Genetics Comprehensive Eiplepsy and Seizure Panel (test code: 7347); https://www.preventiongenetics.com/testInfo?val=PGmaxTM-%252D-Comprehensive-Epilepsy-and-Seizure-Panel) (test code: 16005), other epilepsy-related (M.-W. Zhang et al., 2024), autism (SFARI genes; https://gene.sfari.org/database/human-gene/) (Abrahams et al., 2013), ID (SysNDD phenotype search “intellectual disability”, category: definitive; https://sysndd.dbmr.unibe.ch/Phenotypes) (Kochinke et al., 2016), Parkinsons’s disease (Gene4PD, rare genes; http://genemed.tech/gene4pd/download) (Li et al., 2021), cancer (OncoKB; https://www.oncokb.org/cancer-genes) (Chakravarty et al., 2017; Suehnholz et al., 2024), hypertension (CVD Atlas, PedAM Disease-variant association; https://ngdc.cncb.ac.cn/cvd/disease/CVDD000096) (Qian et al., 2024), rheumatoid arthritis (RADB: http://www.bioapp.org/RADB/index.php/Index/index) (R. Zhang et al., 2014). Human disease gene IDs were converted to mouse gene IDs prior to overlap testing based on the Mouse Genome Informatics nomenclature.

### Drug Treatment and Preparation

Experimental drug testing was performed in non-transgenic mice. The HTR1F agonist LY344864 was ordered from Med Chem Express (Cat. No.: HY-13788). LY344864 was reconstituted in DMSO to stock concentration of 60mg/mL, aliquoted, and kept -80C. On the day of use, LY344864 was resuspended as 0.5% DMSO/drug in corn oil. LY344864 was delivered by intraperitoneal injection at 8 mg/kg. Corn oil containing 0.5% DMSO was used as vehicle control. Non-transgenic mice were injected once-daily for 3 days from P28-P30. One hour after the final injection, PTZ was administered, and seizure responses were recorded.

The Let-7 inhibitor was ordered from Integrated DNA Technologies with the sequence AACTATACAACCTACTACCTCA. A non-targeting scrambled sequence ATAACACTCTAACCACTATCAC was used as a control. Each oligonucleotide was fully modified with 2-methoxyethyl bases and a phosphorothioate backbone. 20 nanomoles of Let7 inhibitor, corresponding to 173.4 micrograms, was delivered by intracerebroventricular injection at age 3 months as previously described (DeVos & Miller, 2013). Let7 inhibition in the CNS using the indicated sequence has previously been validated (Dubinsky et al., 2014; Koval et al., 2013). Four weeks after injection, mice were administered PTZ, and seizure responses were recorded.

### Behavioral Statistics

Multiple linear regressions were used to examine association between seizure score with mouse sex or weight. Seizure scores were non-normally distributed ordinal values, therefore non-parametric two-tailed Mann-Whitney tests were used to compare between treatment groups. Seizure latencies were compared between treatment groups using log-rank (Mantel-Cox) test.

### Data and Code Availability

Raw and processed data will be available through GEO (accession pending). Code will be posted on Bitbucket upon publication.

## RESULTS

### Recording pre-seizure Calling Cards profiles in a pharmacological model of acute seizures

Studies in mice have shown that there is substantial variability in seizure susceptibility across genetically-identical individuals (Aydin-Abidin et al., 2011; Copping et al., 2019; DeVos et al., 2013; Yuskaitis et al., 2021). Therefore, we sought to record pre-existing variation in enhancer activity and correlate inter-individual difference in activity with seizure severity in a PTZ-induced acute seizure model. We delivered conditionally expressed (FLEX, Front Flip), AAV calling cards viral vectors (**Fig 1A**) to the cortices of a cohort of P0-1 mouse pups of the Syn1::Cre genotype. These CC reagents record the binding of BRD4, an epigenetic factor which marks active enhancer regions (Cammack et al., 2020; Gogol-Döring et al., 2016; Latif et al., 2021). The FLEX system ensures recording occurs in neurons and limits the background contribution from other cells. At P28, after recording pre-seizure CC occupancy profiles in each animal, mice were administered acute seizures through intraperitoneal injection of the GABA-A receptor antagonist PTZ at 65 mg/kg. Mice were observed for 15 minutes and were then immediately euthanized, reasoning 15 minutes would allow insufficient time for any seizure-related CC insertions and expression (especially relative to the 4 weeks of recording that have already occurred), while allowing long enough to stratify seizure responses. Cortical tissues were then collected for CC analysis (**Fig 2A**). Seizures were scored based on a modified Racine scale, ranging from 1 (immobility) to 5 (generalized tonic-clonic seizure) to 8 (death). We therefore trichotomized animals by seizure scores into mild (score 1-4), moderate (score 5), or severe (score 6-8) (**Table 1**). Of the 32 mice tested, 12 were mild responders, 9 were moderate, and 11 were severe (**Fig 2B**). Multiple regression analysis revealed that seizure scores were not related to factors such as sex or weight, likely due to our controlling for weight with dosing (**Fig S1A,B**).

**Figure 2.**
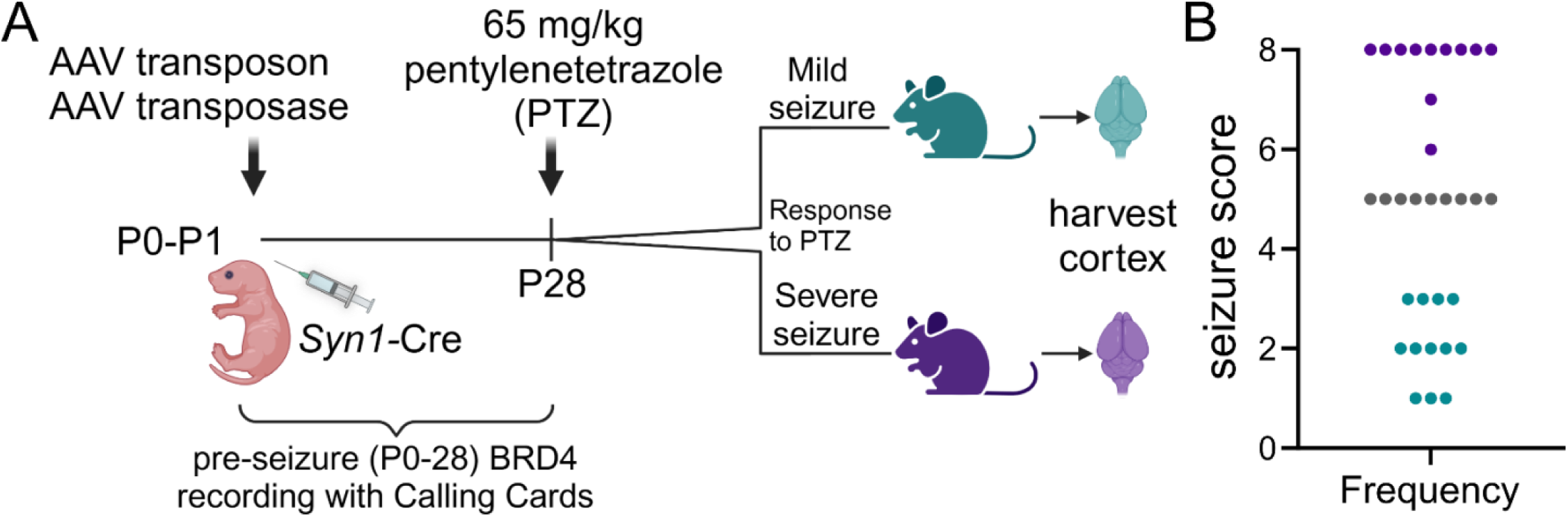
Mice exhibit variable seizure responses to PTZ administration. (A) Experimental paradigm of recording molecular states from P0 to P28 prior to seizure with Calling Cards. (B) Frequencies of seizure score by modified Racine scale after pentylenetetrazol (PTZ) injection. Categorization for seizure severity into mild, moderate, or severe responses are indicated by cyan, grey, or magenta respectively.

### Calling Cards identifies enhancer loci related to later seizure responses

To test whether epigenetic enhancer activity (indicated by BRD4-directed CC insertions) might predict seizure severity, we sought to compare CC profiles between mice that experienced mild or severe responses. Therefore, we prepared and sequenced CC libraries from all mild (12) and severe (11) responders to recover CC insertions. To increase our sensitivity to detect unique insertions, libraries were prepared from separate tissue fragments of each cortex (up to six per animal). We selected the cortex for analysis because the cortex is quickly activated following intraperitoneal PTZ administration in rodents (Keogh et al., 2005). Animals passing QC metrics (methods) were included for downstream analysis, leaving 10 mice per group. CC enhancer profiles were generally very similar across animals within each group (**Fig S2A,B**). As expected, we found that insertions were distributed across the genome (**Fig S2C**) and could not be attributed to any one mouse (**Fig S2D**). We used peak calling to define regions of enriched insertion density, for a total of 16,872 peaks. We expected that these peaks should include known enhancer sites. Indeed, peaks were located predominantly within intergenic regions and introns, rather than promoters, as would be expected (**Fig S2E**). Furthermore, overlaying CC peaks with ChIP-seq datasets of histone modifications from developing mouse forebrain (He et al., 2020) revealed strong enrichment of active enhancer-associated marks H3K27ac (**Fig 3A,B**) and H3K4me1 (**Fig S2F**), and de-enrichment of repressive mark H3K27me3 (**Fig S2G**), as expected from prior work establishing the AAV CC system (Cammack et al., 2020).

**Fig 3.**
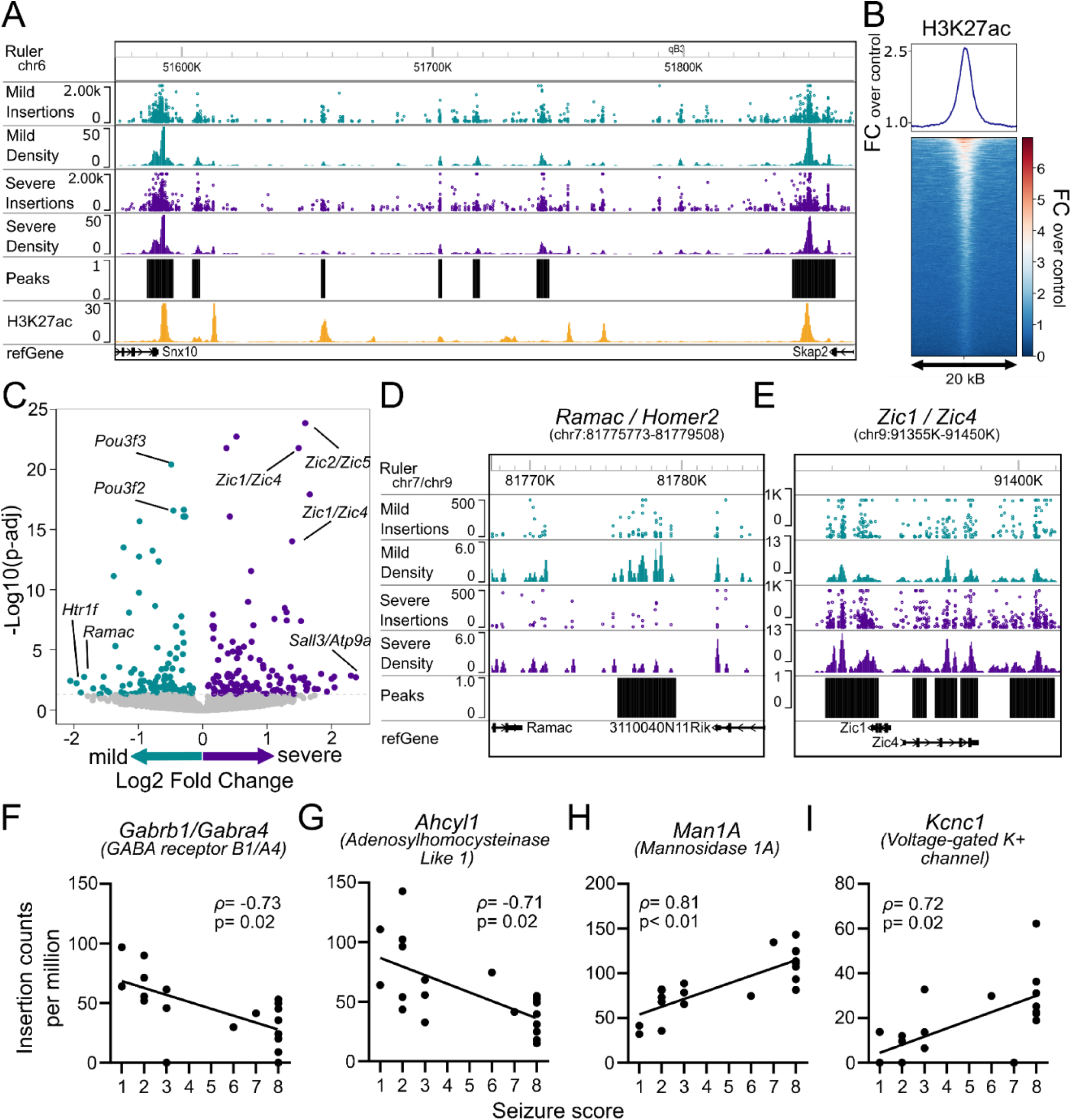
Enhancer usage near hundreds of genes associates with seizure severity. (A) Representative genome browser track depicting number of insertions, insertion density, peaks annotated by CC pipeline, and H3K27ac ChIP-seq density from P0 mouse forebrain (ENCODE) around the *Snx10* and *Skap2* loci (chr6: 51500K-51900K). Insertions enriched in mild or severe responses are separated, and annotated peaks represent genomic locations with strong density across responders. The H3K27 track demonstrates a presence of established enhancer sites near peaks. (B) Enrichment plot (top) and heatmap (bottom) of H3K27ac ChIP-seq (same as A) relative enrichment, quantified as fold change over ChIP-seq control, centered on CC peaks. (C) Volcano plot depicting log2 fold change of differential peaks between mild (green) and severe responders (magenta) with mild enrichment represented by negative fold changes; and severe with positive fold change. Peaks are annotated with nearby genes. (D) Genome browser track for representative peak around *Ramac/Homer2* (chr7:81775773-81779508) depicting differential enrichment of mild insertions. (E) Genome browser track for multiple representative peaks around *Zic1/Zic4* (chr9:91355K-91450K) depicting differential enrichment of severe insertions. (F-I) Spearman correlations between individual animal seizure scores and insertion counts at loci near (F) *Gabrb1/Gabra4* (Spearman ρ= -0.73, p=0.02), (G) *Ahcyl1* (ρ= -0.71, p=0.02), (H) *Man1a* (ρ= 0.81, p<0.01), and (I) *Kcnc1* (ρ= 0.72, p=0.02).

To test the hypothesis that there might be loci whose developmental epigenetic activity predicts later seizure presentation, we compared the number of insertions falling within each peak between mild and severe responders. We observed 110 regions enriched for CC binding in mild-responding animals and 133 for severe responders (**Fig 3C** and **Table 2**). To ensure these differential peaks represented robust epigenetic differences related to seizure vulnerability and not mere individual differences, we shuffled the comparison groups 20 times to comprise animals with mixed seizure severity scores, and performed the same peak calling and differential analysis. Indeed, the proportion of differential peaks we observed in our dataset (1.44%) was significantly higher than that of any of the shuffled arrangements (mean 0.51% , st. dev. 0.35%, p < 0.05), indicating there is a biological coherence to these group assignments that was better than chance (**Fig S2H**).

We next annotated peaks with nearby genes and plotted the directionality of peaks toward mild or severe responders. As an example, we found relative enrichment in CC insertions around the *Ramac/Homer2* gene in mild animals (**Fig 3D**). Similarly, we observed heavy enrichment of CC insertions in peaks around *Zic1/Zic4* in the severe-response mice, with seven peaks reaching significance (**Fig 3E**). Among the strongest enrichment in mild responders occurred around *Htr1f* and *Heatr3*, and *Sall3* and *Bche* for severe. We found enrichment near both coding and non-coding genes, such as the microRNA *MirLet7c-1*. Next, we conducted a posthoc analysis on these peaks showing categorical differences to examine the reproducibility across animals and examine the quantitative relationship between severity score and insertion number. Overall, within the set of mild or severe-related peaks, we found that 13 genes were significantly correlated with seizure score. For example, insertions near *Gabrb1/Gabra4* and *Ahcyl1* were negatively correlated with seizure score (**Fig 3F,G**); insertions near *Kcnc1* and *Man1A* were positively correlated with score (**Fig 3H,I**). These results indicate that CC occupancy at these peaks (BRD4 sites) is predictive of seizure outcomes.

### Genes associated with seizure responses are associated with neurodevelopment and epilepsy

If pre-existing variation in neuronal enhancer activity predicts seizure severity, then genes near differentially active enhancers should be enriched for neurodevelopmental pathways and genes previously implicated in seizure disorders. We performed Gene Ontology (GO) for mild-response genes or severe-response genes against a background list consisting of all genes annotated near CC peaks. (We note that because only Syn-Cre positive cells (neurons) had transposase activity, all loci should be from neurons). Surprisingly, even against this background of neuron loci, we found enrichment of multiple neurodevelopmental pathways in the severe - response genes, but not in mild-response genes. For example, enriched processes associated with severe genes included “generation of neurons” and “neuron differentiation” (**Fig 4A**). Interestingly, severe-associated genes were also enriched for “L-glutamate transmembrane transporter activity,” such as transporters (*Slc1a7*, *Slc7a11)* and receptors (*Grm1*) (**Fig 4A**). These suggest severe responders might carry a greater proportion of neurons in a more neurodevelopmentally immature state and/or might have cell populations more polarized towards excitatory neurons.

**Figure 4.**
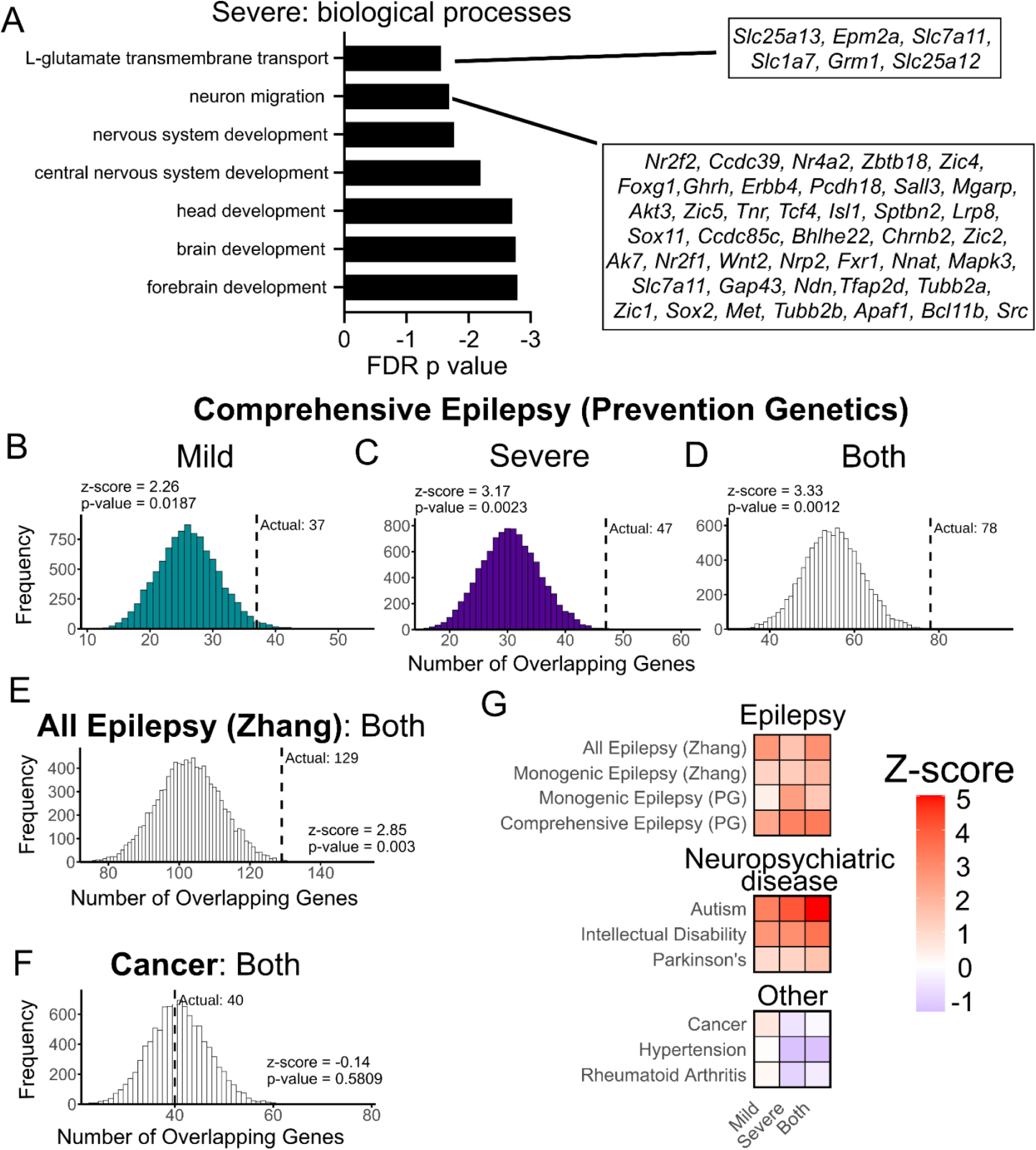
Seizure response-related genes are enriched in neurodevelopmental genes and linked with human risk for seizure disorders. (A) Gene Ontology enrichment for severe genes. Representative genes are annotated for “L-glutamate transmembrane transport” and “neuron migration” processes. (B-D) Distribution of expected gene overlaps between CC genes identified for mild (B), severe (C), or both (D) and comprehensive epilepsy/disease genes list from Prevention Genetics (PG). Observed overlap between significant gene sets indicated by black dashed line. (E) Distribution of expected gene overlaps between CC genes identified for both response groups (mild and severe) and comprehensive epilepsy/disease genes list from Zhang et al 2024. (F) Distribution of expected gene overlaps between CC genes identified for both response groups and gene list associated with cancer. (G) Heatmap summary of z-scores for overlap between mild and severe gene sets and disease genes.

We were specifically interested in whether genes from either list were enriched with established epilepsy genes, either related to monogenic forms of epilepsy or identified as a risk factor in common variant genome wide association studies. This would be consistent with a model where slight epigenetic variations in activities across numerous established epilepsy genes could summate to alter risk. To measure enrichment for known epilepsy genes, we tested our mild and severe gene sets against all genes identified by CC peaks using permutation testing. As a negative control, we included lists of genes implicated in cancer, hypertension, and rheumatoid arthritis. Prevention Genetics clinical panels include a list of seizure and epilepsy genes for which seizures are a major clinical feature (**Fig S3G-I**), as well as a comprehensive list of seizure-and epilepsy-associated genes (**Fig 4B-D**). We found that of the 326 mild genes, 8 were associated with monogenic epilepsy (z-score 0.42) (**Fig S3G**), and 37 with broader epilepsy associations (z-score 2.26) (**Fig 4B)**. Among the 381 severe genes, 15 were associated with monogenic epilepsy (z-score 2.48) (**Fig S3H**), and 47 with broader epilepsy associations (z-score 3.17) (**Fig 4C)**. A recently updated list of seizure genes (M.-W. Zhang et al., 2024) includes and categorizes genes related to epilepsy with varying levels of clinical evidence, ranging from monogenic forms of epilepsy to potentially-associated genes (**Fig S3A-F**, **Fig 4E)**. Of the mild genes, 5 overlapped with monogenic epilepsy genes (z-score 1.17) (**Fig S3D**). Of the severe genes, 6 overlapped with monogenic epilepsy genes (z-score 1.35) (**Fig S3E**). In total, 129 genes from both mild and severe groups (686) overlapped with the list of all epilepsy genes (z-score 2.85) (**Fig 4E**). Owing to a similar etiology, we expected to observe overlap with other neurodevelopmental disorders. Indeed, mild and severe genes each showed strong overlap with autism genes (Abrahams et al., 2013) (35 for mild, z-score 3.2; 44 for severe, z-score 4.08) (**Fig S3J-L**) and intellectual disability (Kochinke et al., 2016) (44 for mild, z-score 2.71; 51 for severe, z-score 2.89) (**Fig S3M-O**). As expected, the overlap with cancer genes (Chakravarty et al., 2017; Suehnholz et al., 2024)(22 mild, z-score 0.62; 20 severe, z-score -0.59) and other less related diseases (hypertension (Qian et al., 2024), rheumatoid arthritis(R. Zhang et al., 2014)) across both response groups was not significantly different from other genes nearby to CC peaks (**Fig 4F, Fig S4)**. Even the overlap with Parkinson’s disease genes (Li et al., 2021) (5 mild, z-score 0.97; 6 severe, z-score 1.13) did not significantly differ from that of other CC genes (**Fig S3P-R**), despite its being a neurological disease, potentially due to its neurodegenerative rather than neurodevelopmental etiology. Overall, our analysis indicates there is significant overlap between the epigenetic risk loci defined here and previously defined genetic risk loci.

### Manipulation of seizure-response genes modifies seizure severity or latency

While there was significant overlap with known epilepsy genes, most CC implicated loci did not contain known epilepsy genes. Therefore, we predicted that these new seizure response-related genes might represent novel pathways influencing seizure activity, and further, targetable pathways for future therapeutic leads. We screened the list of target genes for those that had small molecule antagonists or agonists using the Gene Drug Interactive Database. We found 56 such genes that had an interaction score of at least 10, representing likely target engagement (**Supplemental Table 1**). Owing to the availability of highly-selective, blood-brain barrier penetrant agonists, we selected *Htr1f*, which encodes the serotonin receptor 1f, for further study. Additionally, we had previously developed an antisense oligonucleotide (ASO) inhibitor for another potential target non-coding microRNA *Mirlet7c-1* that can be delivered safely to the central nervous system. A peak at *Htr1f* was enriched in mild responders (mild/severe log2FC of 1.96), and two peaks at *Mirlet7c-1* were modestly enriched in severe responders (log2FC 0.59 and 0.16; **Fig 3C, Fig S6**, and **Table 2**). Notably, neither HTR1F or Let-7 has any existing ties to epilepsy disorders.

*Htr1f* encodes the serotonin receptor 1F, which is an inhibitory G-protein coupled receptor, and its most abundant expression in the central nervous system is in the retina and cortex (Acevedo-Triana et al., 2017). To evaluate whether HTR1F activity can influence seizure responses, we delivered to P28 mice either vehicle or 8 mg/kg HTR1F agonist LY344864 by intraperitoneal injection for 3 days, then administered PTZ (**Fig 5A**). LY344864 is a highly selective serotonin receptor 1F agonist that can cross the blood-brain-barrier (Phebus et al., 1997; Scholpa et al., 2018). We evaluated both the maximal seizure score and the latencies for each mouse to achieve a given score. LY344864 did not influence seizure score, but significantly extended time until mice experienced seizure responses (**Fig 5B**). The first response, usually represented by freezing or immobility behavior, was delayed by 25 seconds (median) (**Fig 5C**). Score 3, which is marked by a “tail pop”, trended to a delay of 30 seconds (**Fig 5D**). Score 5, which is the onset of a generalized tonic-clonic seizure, was delayed by 180 seconds (**Fig 5E**).

**Figure 5.**
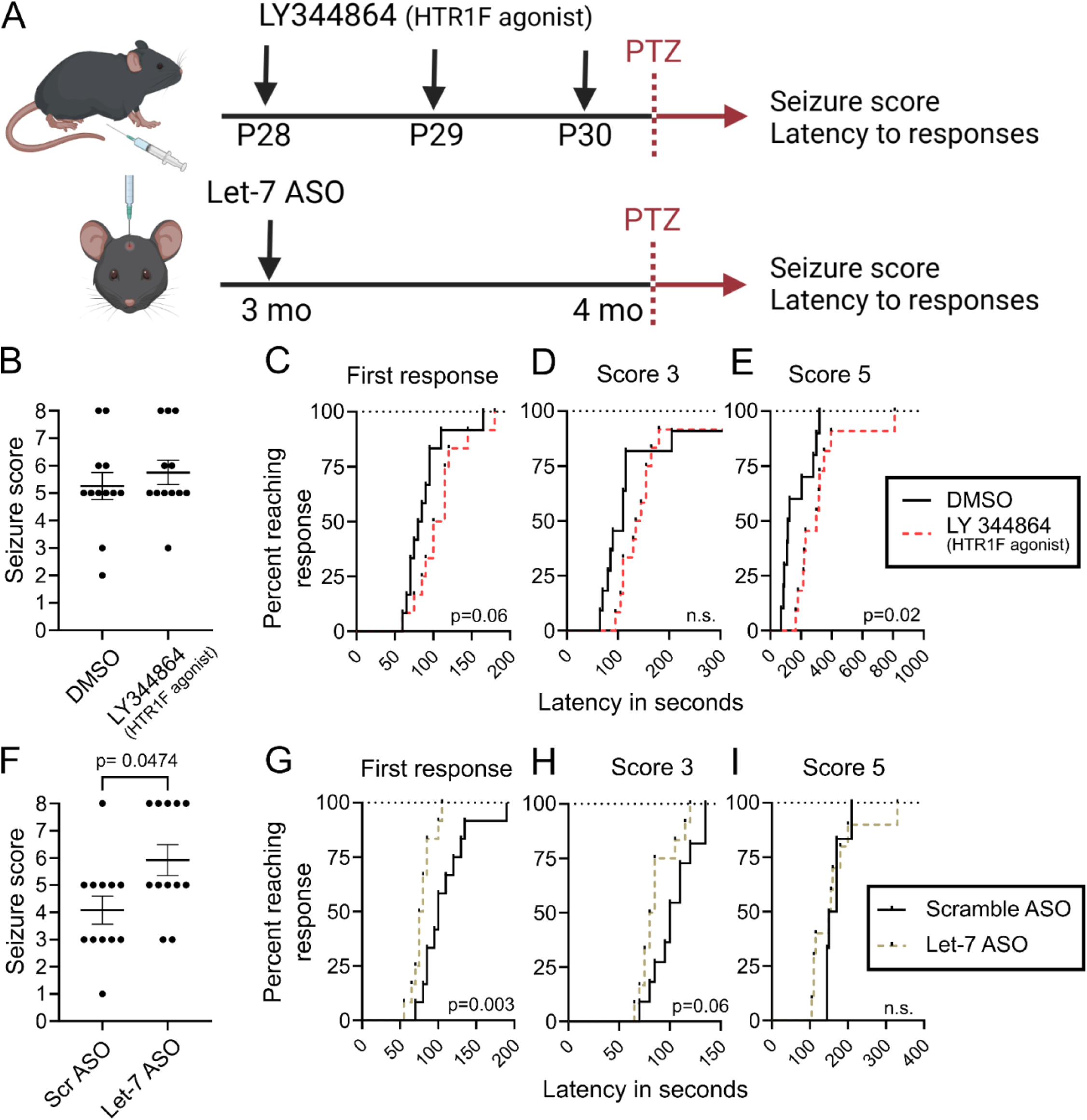
Pharmacological manipulation of seizure associated genes influences seizure responses. (A) Paradigm for treatment of mice with HTR1F agonist LY344864 or Let-7 antagonist (antisense oligonucleotide [ASO]) and subsequent administration of PTZ to characterize seizure responses. (B) Mice were treated with HTR1F agonist or DMSO vehicle for 3 days at P28 by daily intraperitoneal injection and responses to seizure induction were characterized. Seizure scores were plotted between control treatment versus LY344864 (n=12,12). (C-E) Cumulative distribution demonstrating time in seconds between administration of PTZ and onset of (C) first response (n=12, 12; 2-way ANOVA; p=0.06), (D) score 3 (n=11,12), (E) or score 5 (n=10, 11; p=0.02). (F) Mice were treated with Let-7 ASO for 4 weeks from 3 months of age by intracerebroventricular injection, and responses to seizure induction were characterized. Seizure scores were plotted between scramble ASO sequence and Let-7 ASO (n=12,12; p=0.0474). (G-I) Survival curves demonstrating time in seconds between administration of PTZ and onset of (G) first response (n=12, 12; 2-way ANOVA, p=0.003), (H) score 3 (n=11, 12; p=0.06), (I) or score 5 (n=6,10).

Let-7c is a member of the Let-7 miRNA family, which share an identical seed sequence for target binding. Let-7 miRNAs are widely-expressed throughout the brain and suppress hundreds of target mRNAs, including those influencing neuronal synaptic development (McGowan et al., 2018). To evaluate whether Let-7 inhibition could influence seizures, we delivered an ASO to neutralize Let-7 miRNAs by intracerebroventricular injection to 3 month old mice, an age at which ASOs can be administered ICV easily (**Fig 5A**). After one month, we administered PTZ and recorded seizure responses. Let-7 inhibition worsened seizure score and hastened the first response by 22.5 seconds with a trend to delaying score 3 by 17.5 seconds (**Fig 5F-I**).

## DISCUSSION

CC provides a unique opportunity to record epigenetic activity for future readout. Using CC, we sought to evaluate if PTZ-seizure severity can be predicted by antecedent molecular states. We found activity at 243 genomic loci that distinguished mild from seizure responders, and upon mapping these sites to nearby genes, we observed an enrichment with neurodevelopmental pathways and epilepsy risk genes. Finally, we demonstrated the utility of this strategy to identify novel seizure-related targets by pharmacologically manipulating activity of two genes in these loci, HTR1F and Let-7c. Our results support the hypothesis that epigenetic risk factors during neurodevelopment can influence seizure vulnerability, and broadly that CC may be used to probe antecedent events related to non-genetic risk factors for a variety of outcomes.

Our data suggest that natural variation in epilepsy risk can arise from stochastic trajectories that are enacted postnatally and that are tuned across early development. However, the term ‘epigenetics’ classically encompasses a range of non-genetic influences. For example, one could hypothesize differences in either cell proportions or in cell states could explain some of the observed CC differences related to seizure vulnerability. Consistent with the first hypothesis, we found enrichment with specific neuronal markers, such as *Bcl11b (Citp2)* and *Sox2* in severe and *Pou3f3 (Brn1)* and *Pou3f2 (Brn2)* in mild responders (Loo et al., 2019). This observation hints that differences in proportions of cells, such as in numbers of *Brn1/Brn2*+ (Layer 2-4 excitatory neurons) or *Ctip2*+ (Layer 5 excitatory) neurons, arising during development could be a contributing factor to seizure responses. Another example was the enrichment of “glutamate signaling” in GO for severe -response genes. This might suggest greater abundance of excitatory neurons, whereas the enrichment of insertions near GABA receptor subunits (*Gabra4, Gabrb1, Kcdt16*) and transporters (*Slc6a1* and *Slc6a11*) in the mild group might suggest enrichment with inhibitory neurons. On the other hand, epigenetic differences could also reflect cellular states, circuit wiring, or of course, the more recent definition of epigenetics and meaning chromatin states. Indeed, the presence of GABA receptor subunits could reflect differences of excitability in each cell, rather than differences in cell proportions.

The precise timing of when the differential enhancer usage occurred also remains elusive. Because CC records continuously from delivery (P0-P1) (Cammack et al., 2020), epigenetic differences could either be attributed to trajectory over development or the matured state at sacrifice. Even within development, our recording window from P0 to P28 covers the periods of synaptogenesis, synaptic pruning, gliogenesis, programmed cell death, and myelination (Chini & Hanganu-Opatz, 2021; Thion & Garel, 2017; Zeiss, 2021) --any of which could have influenced seizure vulnerability. These questions motivate the need for inducible CC systems to enable temporal control of recording during desired windows. Such systems would also allow for examination of non-genetic factors contributing to a range of phenotypes presenting on longer time scales.

Nonetheless, we found interesting loci that may act epigenetically to influence seizure propensity. Most known genetic forms of epilepsy are related to gain or loss of function mutations in a single gene. We predicted that slight differences in activity of these genes might be associated with increased epilepsy risk. Indeed, mild responses were linked with CC activity near epilepsy genes like *Gabrb1* and *Slc6a1*, for example. Severe responses had greater CC activity near genes such as *Chd2* and *Kcnc1*. Compared to established, clinically-used panels of epilepsy genes (Prevention Genetics and Zhang et al., 2024), we found strong enrichment across both mild and severe genes. This suggests the epigenetic signatures detected by CC might act in either direction; indeed, BRD4 can act as either a repressor or activator, depending on the context (Latif et al., 2021; Liu et al., 2022; Sakamaki et al., 2017).

Polygenic risk for epilepsy has been explored in GWAS mega-analyses from The International League Against Epilepsy Consortium on Complex Epilepsies, and we might expect the CC-tagged risk might also overlap with such common genetic risk. Genome-wide mega-analyses in 2018 and 2023 have respectively identified 16 and 26 risk loci related to various epilepsy subtypes (Abou-Khalil et al., 2018; Stevelink et al., 2023). Of the five loci associated with all forms of epilepsy in their analyses, three were linked with seizure response in our analysis: *VRK2/FANCL* (mapped to *BCL11A*, severe), *RORB* (mild), and *HEATR3* (mild). Of those linked with generalized epilepsy, six sites associated with seizure response: *POU3F3, PTPRK, PCDH7, COX7B2/GABRA4* (mapped to *GABRA2*), *GRIK1*, and *ACVRL1/ACVRL1B* (mapped to *SCN8A*). The locus including *VRK2/FANCL/BCL11A* has been strongly associated with epilepsy risk, and its mapping to *VRK2* or *BCL11A* has not been clear (“Genetic Determinants of Common Epilepsies,” 2014; Stevelink et al., 2023). An interesting observation in our results is the identification of their homologs *Vrk1* and *Bcl11b* as both in loci associated with seizure response, despite being in distinct loci. If both can influence severity, this suggests the human locus’s association may be driven by both genes. Collectively, our data argue that slight variations in activity at both established monogenic epilepsy genes and common variant associated loci occur stochastically, and these variations during neurodevelopment might be predictive of future risk.

We also hypothesized that novel genes could suggest unexplored mechanisms contributing to epilepsy, or targets for therapeutic intervention. One intriguing example in our dataset with strong enrichment in severe - responders was the ZIC family (*Zic1*, *Zic2*, *Zic4*, and *Zic5*). The genomic locus that contains *Zic1* and *Zic4* had seven significant peaks, and the region containing *Zic2* and *Zic5* had five peaks. ZICs are zinc-finger proteins and transcriptional activators that direct neural progenitor proliferation in neurodevelopment (Aruga et al., 2002; Murillo et al., 2015). Given their overwhelming presence in our dataset, we suspect ZIC proteins might also represent novel epilepsy genes and anti-seizure targets, but as yet, no pharmacological agent for this family is available. Most current anti-epileptic drugs operate by subduing neuronal excitability, for example by inhibiting sodium channels or augmenting GABAergic signaling (Sills & Rogawski, 2020), so we sought to evaluate other mechanisms contributing to seizure activity. One gene target identified from our dataset that has established pharmacological tools was the serotonin receptor 1F (Htr1f). HTR1F is an inhibitory G-protein-coupled receptor that responds to serotonin but has relatively unknown functions. While agonism of HTR1F with LY344864 produced modest seizure-modifying activity in our assay, we predict that further development may yield a promising anti-epileptic. The non-coding microRNA Let-7 was also an intriguing target, owing to the presence of two severe-associated peaks near the *Mirlet7c-1* genomic locus. We found that inhibiting Let-7 activity worsened seizure responses, validating its contribution to seizure activity. Furthermore, we anticipate that exploring pathways like those influencing ZIC activity could also inspire novel therapies for epilepsy once new tools to manipulate these pathways have been developed. Overall, epigenetic risk factors such as those suggested here might yield novel drug targets or adjuvants for existing therapies providing untapped strategies to affect disease.

Beyond epilepsy, an individual’s outcomes or risk for neuropsychiatric diseases like Alzheimer’s disease or major depression hinge on natural variation. In these circumstances, epigenetic variability might manifest in an individual’s unique presentation, disease course, or response to therapy. In behavioral neuroscience, this variability might be modeled in “learners” or “non-learners” in social operant tasks that might stratify mice based on social motivation (Maloney et al., 2023), or distinguishes mice that respond to social defeat stress with longer term depressive like behavior (vulnerable vs. resilient mice) (Russo et al., 2012). CC provides the opportunity to isolate the epigenetic factors in balance during prodromal states, to reveal what protective pathways support successful therapeutic response, or to identify factors that make genetically identical individual animals respond differently to the same event. This work should serve as a template for applying this strategy broadly to reveal new mechanisms of disease and targets for therapeutic intervention.

## Acknowledgments

The authors thank Harrison Gabel for helpful comments on the manuscript, the DNA Sequencing Innovation Lab, Genome Technology Access Center at the McDonnell Genome Institute (GTAC@MGI), and the High Throughput Computing Facility at the Center for Genome Sciences and Systems Biology. This work was supported by the Hope Center Viral Vectors Core at Washington University School of Medicine. This work was supported by grants from the National Institute of Mental Health (RF1MH117070, RF1MH126723, 1F30MH136688-01A1 to R.D.M., J.D.D., and M.A.G.); National Institute of Aging (F30AG082394-01A1 to B.D.B.)

## Author Contributions

**Project conceptualization:** B.D.B., M.A.G., A.J.C., R.D.M., J.D.D., T.M.M. **Method development, experiments, and data collection:** B.D.B., D.A.G., S.M.M., M.S., T.C. **Formal analysis:** B.D.B., M.A.G., J.G. Figures and **data visualization:** B.D.B., M.A.G., J.G. **Writing-original draft:** B.D.B., M.A.G., R.D.M., J.D.D., T.M.M. **Writing-review and editing:** B.D.B., M.A.G., D.A.G., A.J.C., R.D.M., J.D.D., T.M.M. **Project coordination:** B.D.B., M.A.G., R.D.M., J.D.D., T.M.M. **Funding acquisition:** B.D.B., M.A.G., R.D.M., J.D.D., T.M.M

---

**Supplemental Figure 1.**
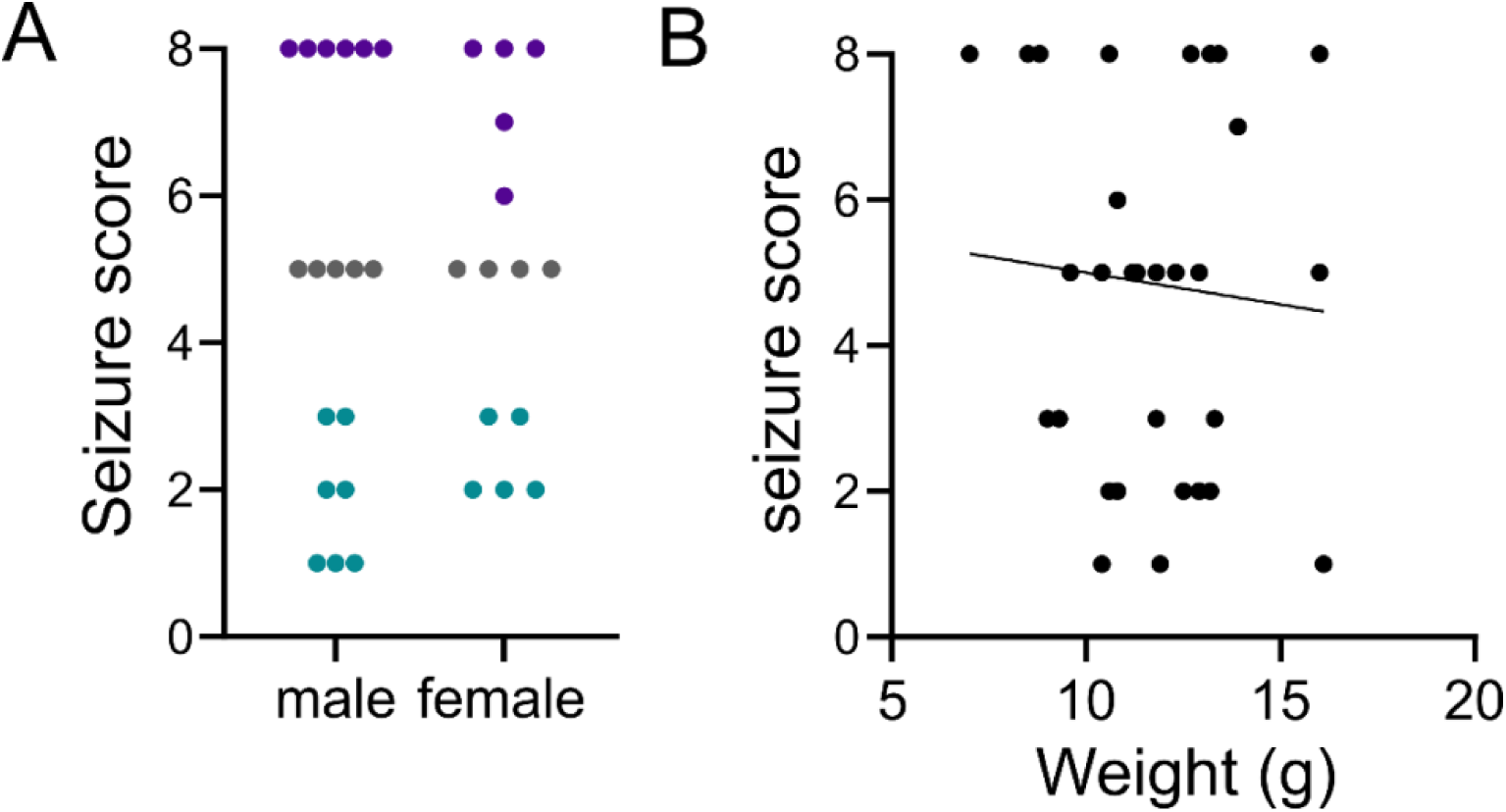
Seizure scores do not vary by sex or weight. (A) Seizure scores were plotted by sex. n = 18 males, 14 females. Student’s t-test p= 0.500. (B) Seizure scores were plotted by weight. n = 32. Pearson R2 = 0.005648, p = 0.6827.

**Supplemental Figure 2.**
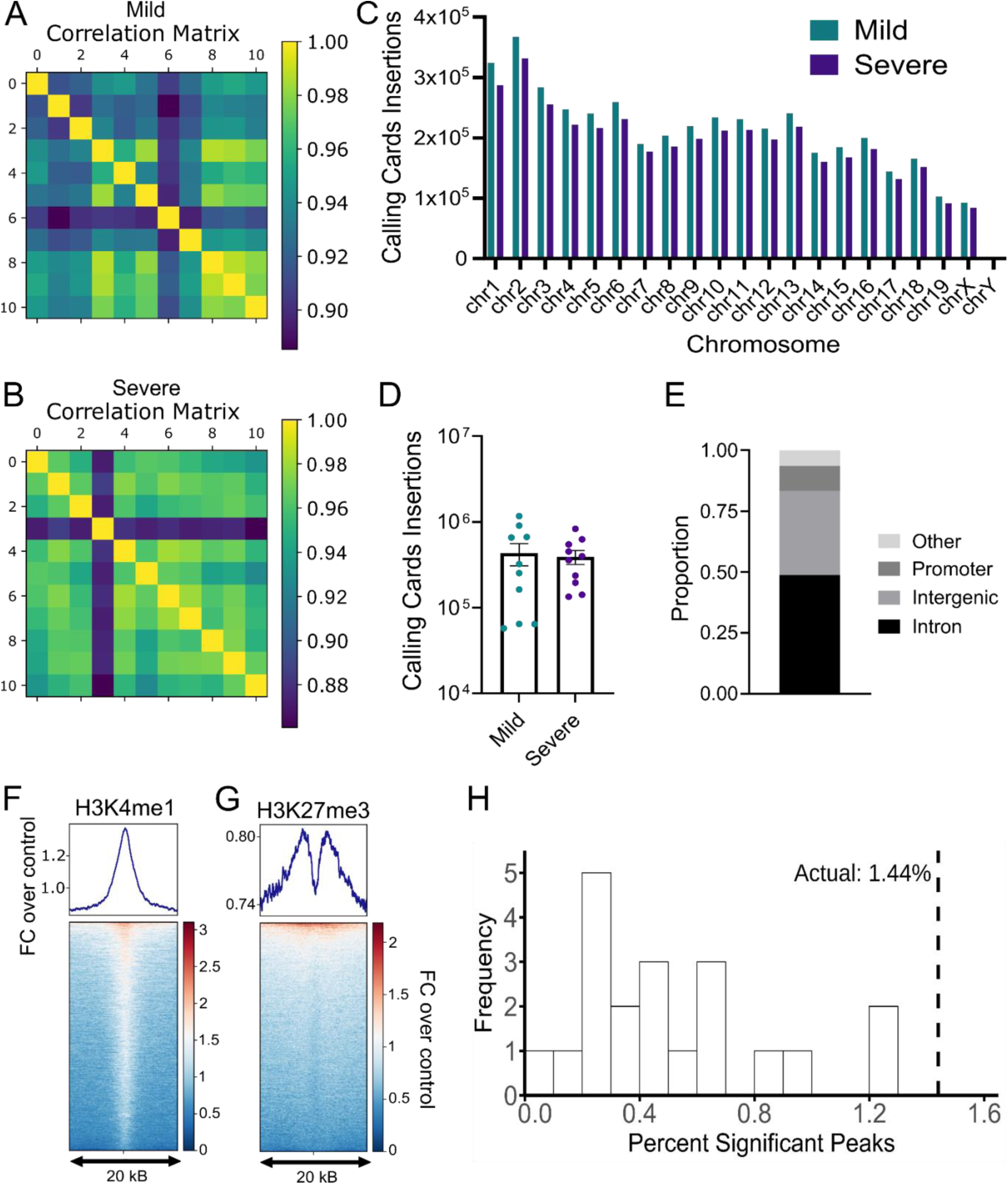
BRD4 Calling Cards targets enhancers across the genome. (A-B) Correlation matrices of insertions per peak by animal for mild (A) and severe (B) responders. (C) Distribution of insertion counts per chromosome in mice with severe (n = 10) and mild (n = 10) seizures. (D) Insertion counts by animal for mild and severe groups. (E) Genomic features of peaks (n = 16,872) called from Calling Cards insertions. “Other” includes exon, 3’ UTR, 5’ UTR, TTS, miRNA, ncRNA, and pseudo. (F,G) Enrichment plot (top) and heatmap (bottom) of CC peaks and H3K4me1 (F) and H3K27me3 (G) ChIP-seq relative enrichment, quantified as fold change over ChIP-seq control, centered on CC peaks. ChIP-seq data was obtained from P0 mouse forebrain (ENCODE). (H) Distribution of differential peak proportions for shuffled insertion profiles (white bars) versus actual dataset (black dashed line).

**Supplemental Figure 3.**
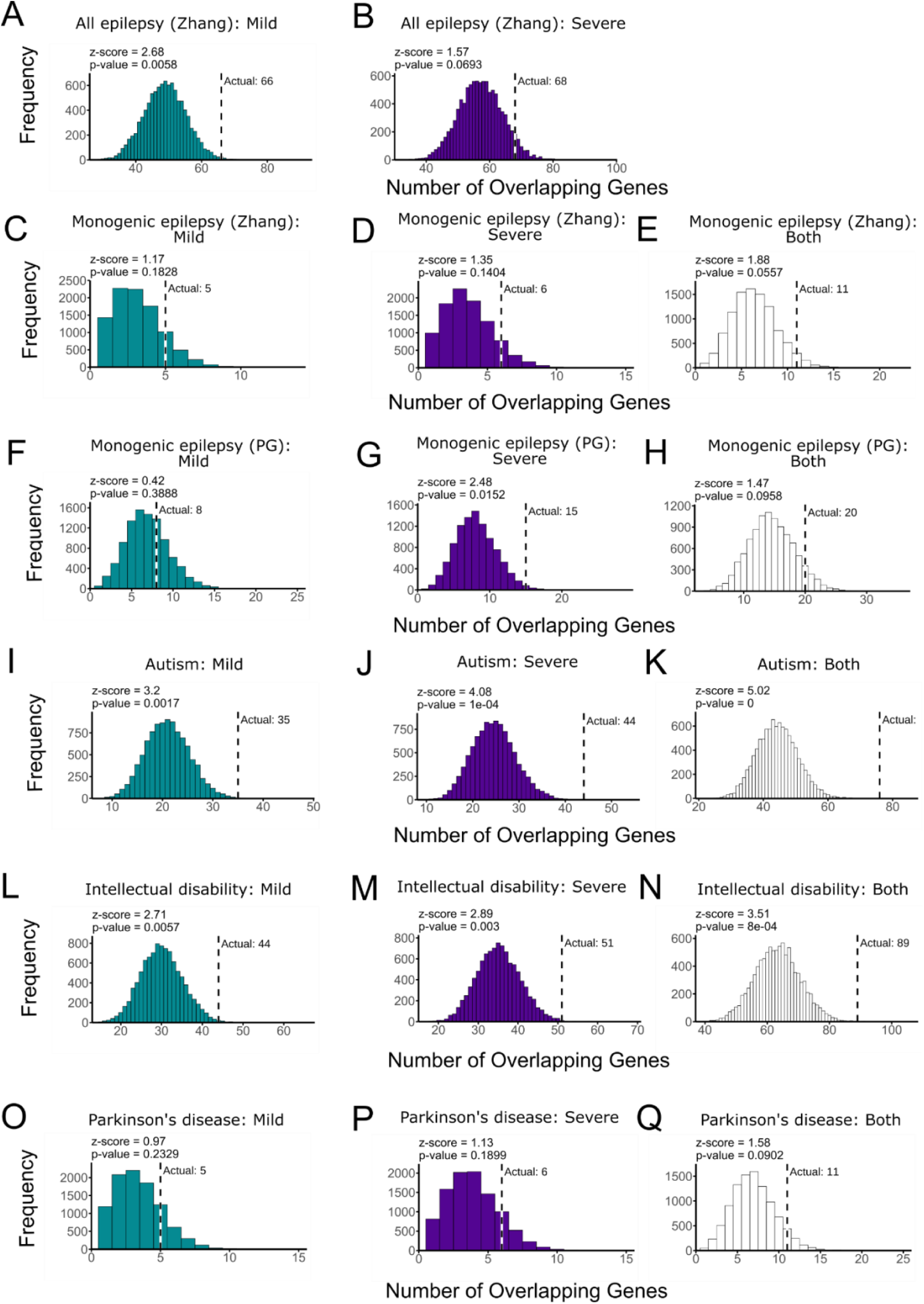
Gene overlaps between seizure response-associated genes with neuropsychiatric conditions. (A-H) Distribution of expected gene overlaps between CC genes identified for mild or severe, or both responses and gene lists associated with (A, B) “All epilepsy” annotated by Zhang et al 2024, (C-E) “monogenic epilepsy” annotated by Zhang et al. 2024, (F-H) “monogenic epilepsy” annotated by Prevention Genetics. (J-R) Distribution of expected gene overlaps between CC genes identified for mild, severe, or both responses and gene list associated with “autism” (I-K), “Intellectual disability” (L-N), or Parkinson’s disease (O-Q).

**Supplemental Figure 4.**
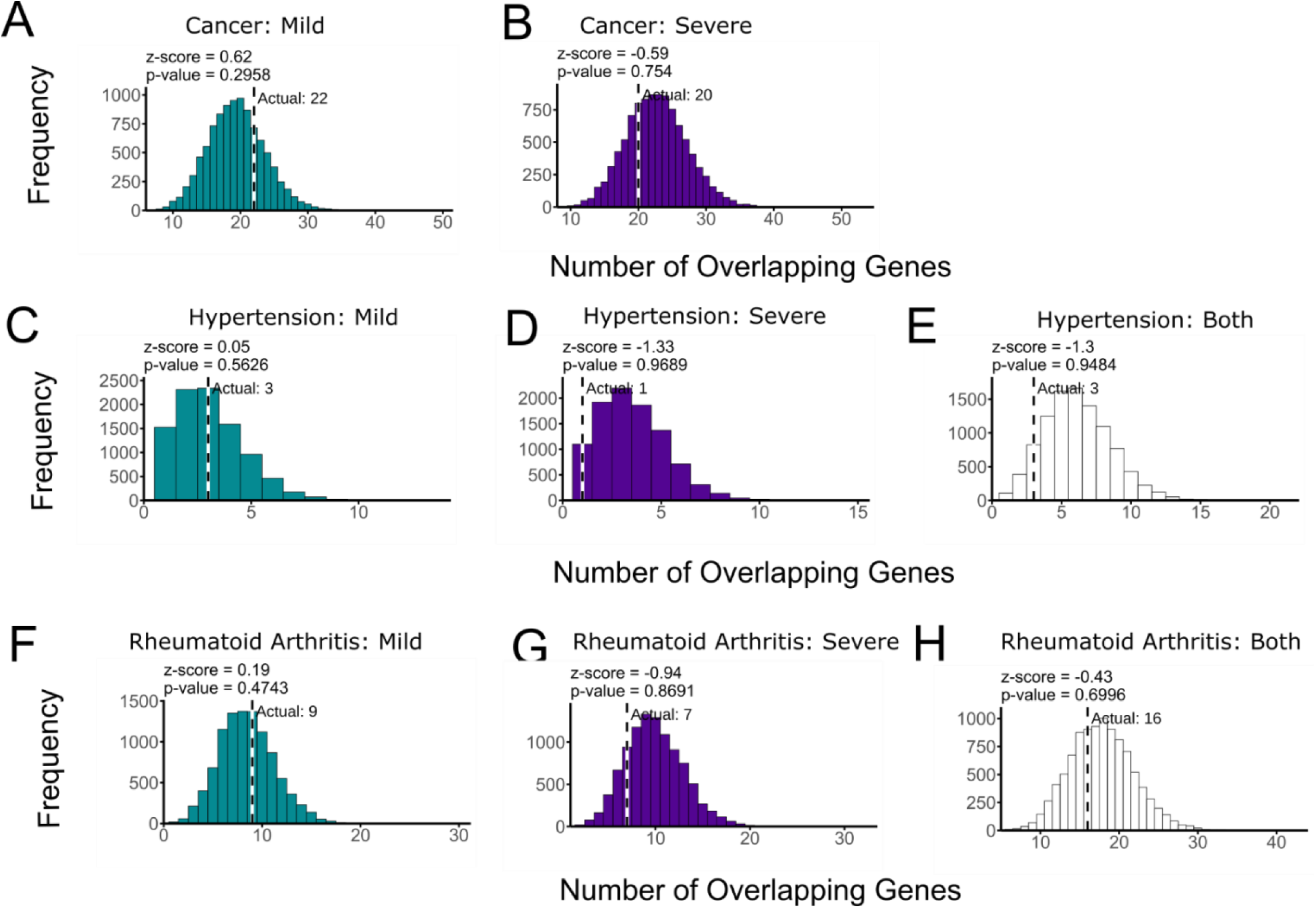
Gene overlaps between seizure response-associated genes with non-neurological conditions. (A, B) Distribution of expected gene overlaps between CC genes identified for mild (A) or severe (B) responses and gene list associated with cancer. (C-E) Distribution of expected gene overlaps between CC genes identified for mild (C), severe (D), or both (E) responses and gene list associated with hypertension. (F-H) Distribution of expected gene overlaps between CC genes identified for mild (F), severe (G), or both (H) responses and gene list associated with rheumatoid arthritis.

**Supplemental Figure 5.**
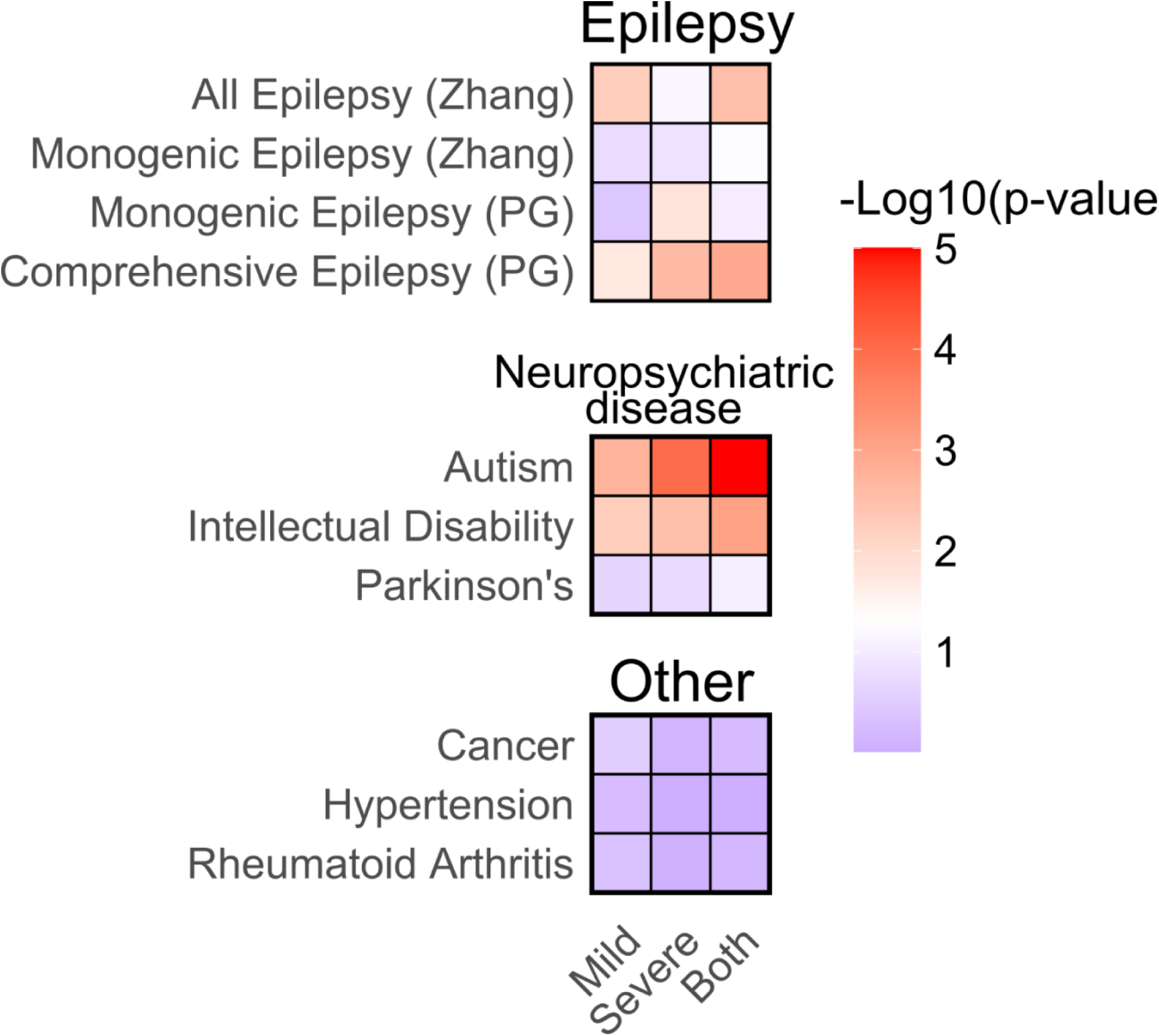
Heatmap depicting p values for overlap analysis between seizure response-associated genes with epilepsy, neuropsychiatric disease, and other gene lists.

**Supplemental Figure 6.**
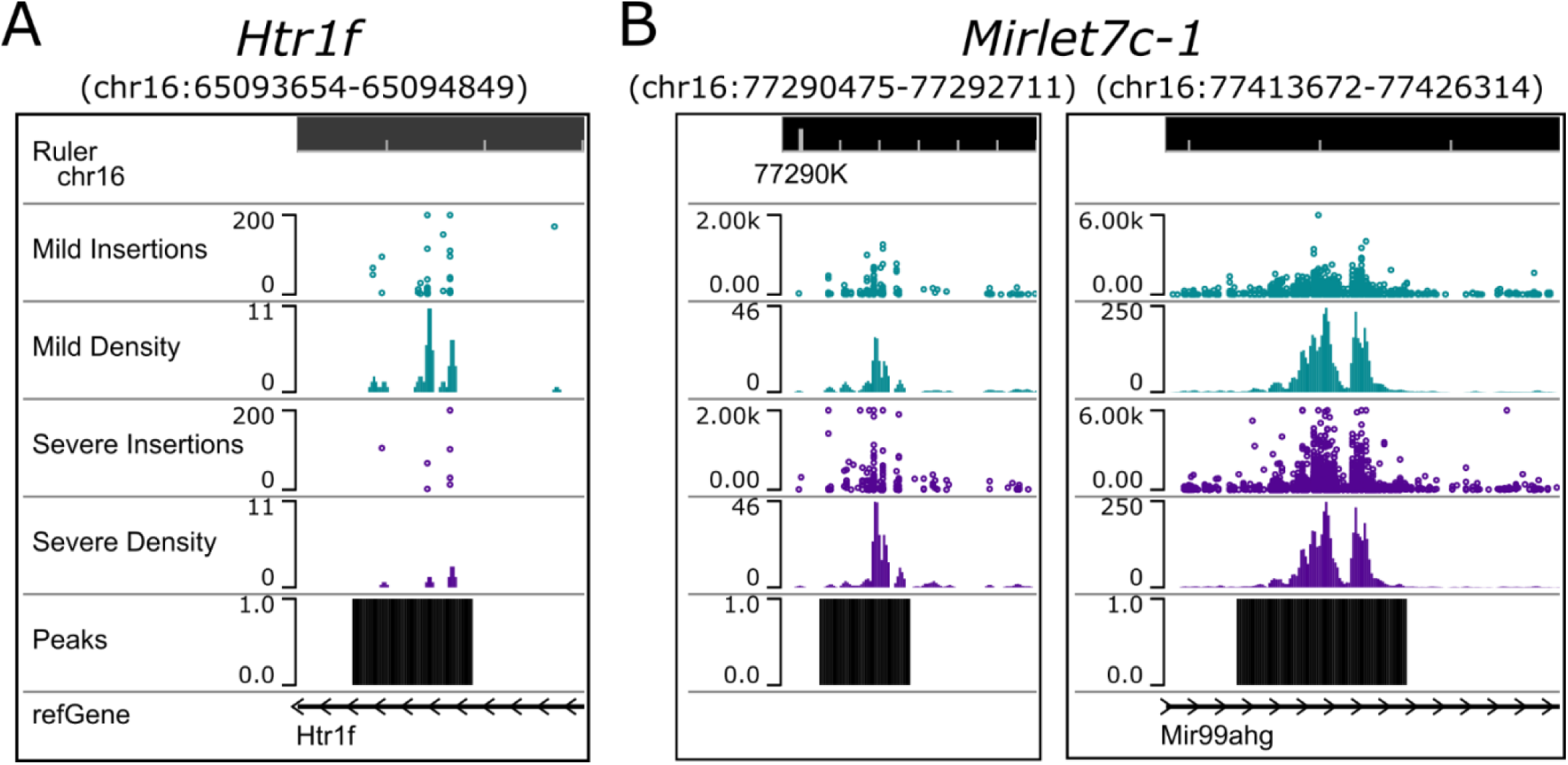
Differential peaks for follow-up studies. Genome browser tracks for peaks around *Htr1f* (A) (chr16:65093654-65094849) and *Mirlet7c-1* (B) (chr16:77290475-77292711 and chr16:77413672-77426314) depicting differential enrichment of insertions.

**Supplemental Table 1. Scoring of gene-drug interactions using dgIDB.** Genes were associated with existing drugs and scoring based on strength of evidence is represented as “interaction score”.

